# Endothelial Function: A Novel Marker to Evaluate the Prognosis of Heart Failure with Reduced Ejection

**DOI:** 10.1101/2025.07.13.664603

**Authors:** Salma Charfeddine, Mohamed Ali Hbaieb, Niez Laribi, Mariem Jabeur, Amine Bahloul, Marwa Jarraya, Hassen Gargouri, Aiman Ghrab, Zied Triki, Tarak Ellouze, Faten Triki, Rania Gargouri, Leila Abid

## Abstract

**Background:** Endothelial function, a key determinant of prognosis in heart failure with reduced ejection fraction (HFrEF), is still frequently under-assessed in clinical practice. The present study aimed to assess endothelial function in patients with HFrEF and investigate its association with echography and hemodynamics over a 3-month medical treatment. Additionally, this study aimed to investigate the association between changes in endothelial function and the incidence of cardiovascular rehospitalizations or deaths.

**Methods:** This prospective longitudinal study included 120 patients with HFrEF. Hemodynamic parameters were assessed using impedance cardiography. Endothelial function was evaluated using digital thermal monitoring to calculate the Endothelial Quality Index (EQI) at baseline and after 3 months. Patients were followed for 12 months.

**Results:** The mean age was 61.9 ± 10.2 years, with a sex ratio of 5:1. 42.5% of patients tend to experience endothelial dysfunction at baseline. After 3 months of optimized therapy, EQI improved significantly (p<0.001), correlating with improved echography and hemodynamic parameters. Over 12 months, there were 5 deaths (4.16%) and 44 heart failure rehospitalizations (36.6%), predominantly among those with severe endothelial dysfunction (p=0.008). Improved EQI was associated with reduced mortality (AUC = 0.82) and rehospitalization risk (AUC = 0.837). A ΔEQI ≥ 0.2 predicted better prognosis (HR: 0.157, 95% CI: 0.054–0.454, p=0.001).

**Conclusion:** Patients with HFrEF exhibited endothelial dysfunction. The improvement in endothelial function after an optimized treatment is associated with an enhancement in echography and hemodynamic parameters. Additionally, endothelial function was a strong prognostic marker.

## Introduction

Heart failure represents a major global public health challenge, with an incidence of approximately 3 cases per 1,000 person-years in Europe (1). According to the latest update of the American Heart Association, heart failure accounts for 9.3% of all deaths in the United States (2).

Chronic heart failure (CHF) is a heterogeneous clinical syndrome characterized by a structural and functional alteration of diastolic and/or systolic function that results in a reduction in cardiac output and or elevated intracardiac pressures with alterations in hemodynamic parameters (3).

Echocardiography is a first-line imaging technique and the reference initial diagnostic tool. It provides essential diagnostic and prognostic information, notably for determining left ventricular ejection fraction (LVEF), estimating filling pressures, and assessing cardiac output.

Additionally, impedance cardiography (IC) is a non-invasive, simple, and cost-effective method used to evaluate cardiac performance by measuring changes in thoracic electrical impedance. It provides hemodynamic parameters correlated with invasive measurements, such as thermodilution.

Despite the progress in the monitoring and evaluation tools, heart failure with reduced ejection fraction (HFrEF) remains associated with impaired quality of life and poor prognosis, marked by high mortality and rehospitalization rates for acute heart failure. Specifically, the 5-year survival rate is only 57.6% (4), 20% of patients are readmitted within the same year, and more than 80% are readmitted within 5 years (5).

Traditionally, follow-up care for patients with HFrEF has focused primarily on cardiac function, with less attention given to vascular function. The endothelium is a single layer of cells lining the blood vessels. Endothelial cells play a crucial role in regulating vascular tone through the synthesis and release of endothelium-derived relaxing factors, including nitric oxide (NO), to induce vasodilatation in response to diverse physiological demands. Furthermore, the endothelium is involved in angiogenesis, hemostasis control, and maintaining blood fluidity by preventing platelet and leukocyte activation (6). Endothelial dysfunction is characterized by an imbalance between vasodilation and vasoconstriction with a reduction in NO production, a redox imbalance, and an increase in the sensitivity to vasoconstrictor substances and pro-inflammatory compounds.

It has been demonstrated that endothelial dysfunction plays a substantial role in the pathogenesis and development of cardiovascular diseases(7). It was advocated that patients with CHF tend to exhibit a lower production of NO and an exacerbation of oxidative stress (8).

Despite its crucial role in predicting prognosis in cardiovascular disease (9), endothelial dysfunction remains largely overlooked in clinical practice, particularly in CHF management.

The present study aimed to assess endothelial function in patients with HFrEF and investigate its association with traditional clinical outcomes (i.e., echography and hemodynamics) over a 3-month medical treatment. Additionally, this study aimed to test the hypothesis that endothelial function could be a prognostic indicator for the long term.

We hypothesize that endothelial dysfunction is a key determinant of heart failure progression and that its assessment could provide valuable prognostic insights while guiding more effective therapeutic strategies.

## Methods

### Participants

Participants included in the present study were adult patients over 18 years old with HFrEF, as defined by the latest 2023 update of the 2021 European Society of Cardiology guidelines (10). The diagnosis was confirmed by echocardiography within the past six months. Patients were excluded if they met the following criteria: 1) worsening heart failure, defined as an emergency hospitalization for acute decompensation within the past week, an emergency department visit in the past week due to worsening symptoms requiring intravenous diuretic therapy, or outpatient treatment involving either intravenous diuretics in an ambulatory setting or an intensification of oral diuretic therapy; 2) kidney disease, indicated by a creatinine clearance <30 mL/min or hyperkalemia >5.5 mmol/L requiring temporary discontinuation of heart failure treatment; 3) invalid endothelial function assessment. Before enrollment in the study, participants were thoroughly informed about the study’s protocol and provided written informed consent. Ethical approval for this study was obtained from the local Ethics and Investigation Committee (Faculty of Medicine of Sfax N° 24/25). This study was conducted according to the Declaration of Helsinki.

### Study design

This longitudinal and prospective study was conducted in the Department of Cardiology at the University Hospital Hedi Chaker Sfax in Tunisia from June 2022 to December 2023.

At the inclusion, an initial consultation was conducted to assess endothelial function and echocardiographic hemodynamic parameters. These parameters were reassessed after 3 months of medical therapy for HFrEF under the European Society of Cardiology recommendations (10). The pharmacological treatment included renin-angiotensin-aldosterone system inhibitors, such as angiotensin-converting enzyme inhibitors (Captopril, Enalapril, or Ramipril), angiotensin II receptor blockers (Valsartan or Candesartan), and angiotensin receptor-neprilysin inhibitors (Sacubitril-Valsartan). Additionally, beta-blockers (Carvedilol, Bisoprolol, or Nebivolol), aldosterone antagonists (Spironolactone, the only prescribed molecule in this category for our study population), and sodium-glucose co-transporter 2 inhibitors (SGLT2) were administered. Patients were monitored for one year after inclusion, with documentation of any major cardiac events, such as hospitalizations for acute heart failure exacerbations or cardiovascular-related deaths.

### Endothelial function

Endothelial function was assessed using an E4-diagnose device (Polymath Company, Tunisia) that fully automates the post-occlusion hyperemia protocol. The E4-diagnose is a non-invasive, high-resolution (0.002°C) skin temperature measuring device composed of two finger temperature sensors, an integrated wrist cuff placed on the dominant forearm, and a portable micro unit controller. Temperature measurements from the non-dominant hand serve as an internal control. A dedicated computer software allowed for data visualization, storage, and export. All measurements were conducted in the morning, in a dimmed and quiet room with an ambient temperature between 22 and 24°C. Patients were required to fast for at least 8 hours, with no smoking and no heavy physical activity for at least 4 hours, and remain seated in a relaxing position at least 20 minutes before the test. To proceed with the measurement, the temperature of the test finger had to exceed 27°C. The protocol included a 5-minute baseline recording period, followed by 5 minutes of cuff occlusion. During occlusion, the temperature of the index finger on the dominant hand decreases due to the interruption of warm blood flow. Upon cuff release, the reactive hyperemic response led to a temperature rebound in the same finger, which was used to evaluate endothelial function via the Endothelium Quality Index (EQI) (11).

Based on the EQI, endothelial function was classified as follows:

EQI > 2: healthy endothelial function

1≤ EQI ≤ 2: mild endothelial dysfunction

EQI < 1: severe endothelial dysfunction

### Echography parameters

Echography parameters were measured at baseline and after 3 months using echocardiography (EPIQ CVx model; Philips, Bothell, WA, USA). The parameters assessed included geometric parameters, left and right ventricular systolic function, myocardial strain, and diastolic filling patterns.

Geometric parameters gather the end-diastolic diameter and the end-systolic diameter. Left ventricular systolic function was evaluated using the biplane Simpson’s method to calculate LVEF. Global longitudinal strain (GLS) was assessed using speckle-tracking echocardiography from standard apical views, providing a sensitive marker of subclinical systolic dysfunction.

Right ventricular systolic function was estimated using Tricuspid Annular Plane Systolic Excursion measured by M-mode imaging at the lateral tricuspid annulus.

Diastolic function was assessed via pulsed-wave Doppler of transmitral flow to measure E and A wave velocities, E/A ratio, and deceleration time. Tissue Doppler imaging was used to measure early diastolic mitral annular velocity (e’), and the E/e’ ratio was calculated as an estimate of left ventricular filling pressures. Furthermore, the average biplane area, left atrial volume, and indexed left atrial volume in end-systole were calculated.

As part of the assessment of filling pressures and pulmonary venous return, the pulmonary artery systolic pressure was estimated from the tricuspid regurgitant jet velocity using the modified Bernoulli equation, with right atrial pressure estimated based on inferior vena cava size and its collapsibility. Elevated pulmonary artery systolic pressure may reflect increased left atrial pressure and advanced diastolic dysfunction, particularly in the setting of post-capillary pulmonary hypertension.

### Impedance cardiography

Hemodynamic parameters were measured using a transthoracic IC (CardioScreen1000, Medis, Ilmenau, Germany). This non-invasive method involves applying two electrodes on the neck and two electrodes on the left side of the chest. The recorded wave reflects the fluctuations in thoracic aortic blood volume and the orientation of erythrocytes as the heart pumps blood from the left ventricle into the aorta. A selection of key parameters is calculated based on the impedance data: flow parameters (i.e., systolic ejection volume and cardiac output), contractility parameters (i.e., the systolic time ratio), fluid parameters including thoracic fluid content, and vascular parameters (i.e., systemic vascular resistance and its index).

## Statistical Analysis

Statistical analyses were performed using the Statistical Package for Social Sciences (SPSS) for Windows (version 26).

Quantitative variables were presented as mean ± standard deviation or as median with interquartile ranges, depending on their distribution. Qualitative variables were expressed as percentages.

The normality of the data distribution was assessed by the Shapiro-Wilk test. The null hypothesis of an absence of differences between groups was tested through a student’s t-test when data were normally distributed, or with a Mann-Whitney U test when they did not fulfill this condition.

The null hypothesis of an absence of difference between groups in categorical variables regarding clinical characteristics was analyzed through the Chi 2 test or the Fisher Test. The null hypothesis of an absence of difference in qualitative variables between the inclusion and 3 months of treatment was tested using the McNemar test.

The null hypothesis of an absence of significant correlation was tested through the Pearson test, as the data met the assumption of a Gaussian distribution.

Multivariate analysis included binary logistic regression to identify independent factors associated with improvement in endothelial function, and Cox proportional hazards regression to identify independent prognostic factors. All variables with a p-value < 0.20 in univariate analysis were included in the multivariate models.

Receiver Operating Characteristic (ROC) curve analysis was used to determine predictive thresholds for mortality and rehospitalization.

An α risk of 5% was retained for all analyses. All statistical analyses were performed with SPSS (Statistical Package for the Social Sciences) version 26 (SPSS Inc., Chicago, IL, USA).

## Results

### Participants

We included in this study 120 HFrEF patients with a mean age of 61.93 ± 10.2 years and a sex ratio of 5:1. Their baseline characteristics were summarized in Table 1.

**Table 1:** Characteristics of the study population at inclusion.

| Characteristics | Mean ± SD or n (%) |
| --- | --- |
| **Cardiovascular risk factors** |  |
| Smoking | 70 (58) |
| High blood pressure | 48 (40) |
| Dyslipidemia | 43 (36) |
| Diabetes mellitus | 51 (43) |
| Family history of coronary artery disease | 79 (66) |
| **Clinical characteristics** |  |
| New York Heart Association class 1 | 16 (13) |
| New York Heart Association class 2 | 39 (33) |
| New York Heart Association class 3 | 41 (34) |
| New York Heart Association class 4 | 24 (20) |
| Systolic blood pressure (mmHg) | 124.4 ±19.18 |
| Diastolic blood pressure (mmHg) | 73.14 ±11.29 |
| **Endothelial function** |  |
| Endothelium Quality Index | 1.27 ±0.6 |
| Healthy endothelial function (EQI>2) | 26 (21.7) |
| Mild endothelial dysfunction (1≤ EQI ≤ 2) | 43 (35.8) |
| Severe endothelial dysfunction (EQI<1) | 51 (42.5) |
| **Echography parameters** |  |
| Left ventricular end-diastolic diameter (mm) | 58.9 ±8.4 |
| Left ventricular end-systolic diameter (mm) | 47 ±9.1 |
| Left ventricle ejection fraction (%) | 29.92 ±7.06 |
| Cardiac output (L/min) | 3.19 ±0.7 |
| Global Left ventricle longitudinal strain (%) | 10.8 ±2.7 |
| Left atrial area (cm<sup>2</sup>) | 23.9 ±5.9 |
| E wave (cm/s) | 83.36 ±25.23 |
| A wave (cm/s) | 56.95 ±26.94 |
| E' wave (cm/s) | 8.2±2.4 |
| Pulmonary artery systolic pressure (mmHg) | 37 ±12.9 |
| S' wave of the right ventricle (cm/sec) | 10.85 ±2.48 |
| Tricuspid annular plane systolic excursion (mm) | 18.27 ±4.07 |
| **Hemodynamic parameters** |  |
| Systolic ejection volume (mL) | 69.14 ±20.5 |
| Cardiac Output (L/min) | 5 ±1.2 |
| Cardiac Index (L/min/m<sup>2</sup>) | 2.73 ±0.6 |
| Systemic Vascular Resistance Index (dyn·s/cm<sup>5</sup>) | 2378.5 ±58.8 |
| Systolic-Time Ratio (1/s) | 0.48 ±0.14 |
| Left Cardiac Work Index (Kg-m/m<sup>2</sup>) | 2.78 ±0.77 |
| Thoracic Fluid Content Index (1/kOhm) | 24.15 ±5.6 |
| **Medication** |  |
| B-Adrenergic receptor antagonist | 86 (71.8) |
| **Aldosterone inhibitor** | 70 (58.9) |
| **Angiotensin-converting enzyme inhibitor** | 63 (53) |
| **Dapagliflozin** | 19 (16.2) |
| **Empagliflozin** | 8 (7.2) |
| **Angiotensin II Receptor Blockers** | 15 (12.7) |
| **Sacubitril-Valsartan** | 6 (5.2) |

**Data are presented as mean ± Standard deviation (SD) or as number (n) and frequency**

### Treatment optimization

At three months, all patients were on optimized treatment, which included the four pillars of therapy for HFrEF. The dosages were either those recommended as optimal or the highest tolerated. The optimal dose of beta-blockers was achieved in 79 (65.8%) patients, while the optimal dose of renin-angiotensin system inhibitors was administered to 68 (56.6%) patients. Additionally, 51 (42.5%) patients received optimal quadruple therapy.

### Effect of 3-month optimized pharmacological treatment

#### Endothelial function

A significant improvement in the EQI was observed after 3 months of optimized treatment, increasing from 1.27 ± 0.62 to 1.44 ± 0.67 (p < 0.001). Additionally, 8 out of the 43 patients initially diagnosed with mild endothelial dysfunction at inclusion were able to restore normal endothelial function by the follow-up visit.

#### Echography parameters

A significant improvement was observed in LVEF, which increased from a mean value of 29.92 ± 7.06% at baseline to 33.98 ± 10.16% after treatment. The paired Student’s t-test revealed a significant mean difference between the pre and post-treatment values (p < 0.001). Similarly, the GLS was increased from a mean value of –10.8 ± 2.7% at baseline to –11.4 ± 3.45% (p = 0.005) after 3 months.

The McNemar test revealed a significant improvement in filling pressures under treatment (p<0.001) (Figure 1). Additionally, normalization of filling pressures after 3 months of optimized treatment was observed in 33 patients who had elevated filling pressures at baseline.

**Figure 1.**
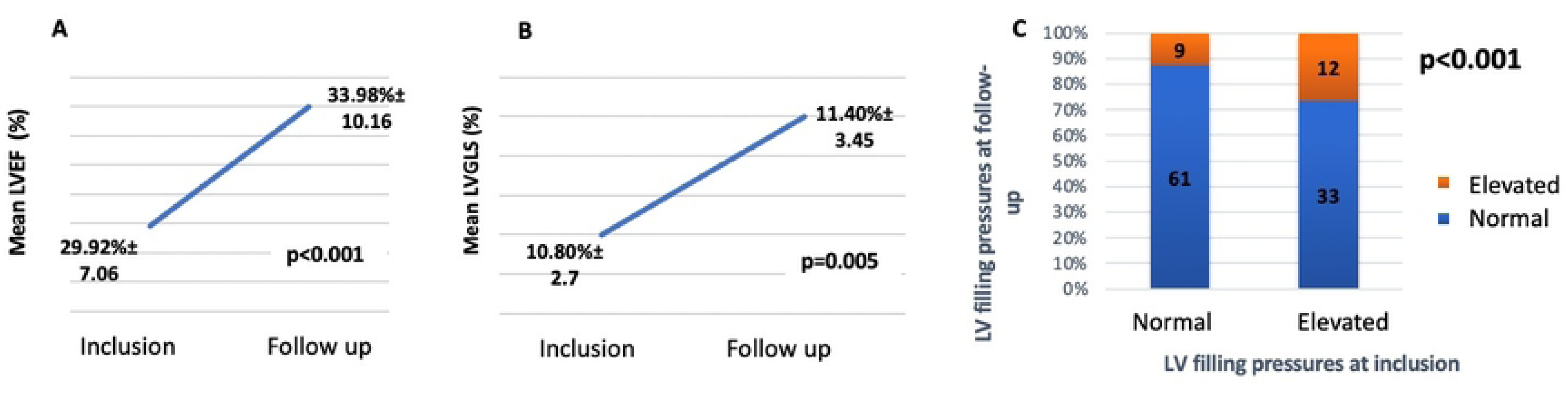
Evolution of the Echocardiography parameters during 3-month follow-up. A. The left ventricle ejection fraction (LVEF), B. The left ventricle global longitudinal strain (LVGLS) and C. The left ventricle filling pressures

#### Hemodynamic parameters

We observed a significant improvement in stroke volume, cardiac output, cardiac index, and cardiac work index. In contrast, there was no significant difference in the systolic time ratio and thoracic fluid content after 3 months of optimized treatment (Table 2).

**Table 2:** Evolution of hemodynamic parameters after an optimized treatment.

|  |  | **Mean± SD** |
| --- | --- | --- |
| **Stroke Volume (mL)** | *Inclusion* | 69.5 ± 21.2 |
|  | *Control* | 81.1 ± 23.7 * |
| **Cardiac Output (L/min)** | *Inclusion* | 4.9 ± 1.1 |
|  | *Control* | 5.8 ± 1.5 * |
| **Cardiac Index (L/min/m<sup>2</sup>)** | *Inclusion* | 2.7 ± 0.6 |
|  | *Control* | 3.2 ± 0.7 * |
| **Systemic Vascular Resistance Index (dyn·s/cm<sup>5</sup>)** | *Inclusion* | 2415.1 ± 589.2 |
|  | *Control* | 2056.5 ± 438.4 * |
| **Systolic-Time Ratio (1/s)** | *Inclusion* | 0.5 ± 0.1 |
|  | *Control* | 0.5 ± 0.1 |
| **Left Cardiac Work Index (Kg-m/m<sup>2</sup>)** | *Inclusion* | 2.8 ± 0.8 |
|  | *Control* | 3.8 ± 2.6 ** |
| **Thoracic Fluid Content Index (1/kOhm)** | *Inclusion* | 24.1 ± 5.6 |
|  | *Control* | 24.3 ± 6.3 |

\*: a significant difference at p<0.05; \*\*: a significant difference at p<0.01

### Factors associated with the improvement of endothelial function after an optimized treatment

Multivariate analysis revealed that treatment with Sacubitril-Valsartan, an optimal dose of beta-adrenergic blockers, as well as guideline-directed quadruple therapy, were predictive factors for the improvement of endothelial function (Table 3).

**Table 3:** Multivariate analysis of factors associated with improvement in endothelial function.

|  | **OR** | **95% CI (Lower)** | **95% CI (Upper)** | **P value** |
| --- | --- | --- | --- | --- |
| **No diabetes** | 2.687 | 1.139 | 6.337 | 0.024 |
| **Statin therapy** | 1.772 | 1.681 | 3.112 | 0.034 |
| **Sacubitril-Valsartan therapy** | 0.367 | 0.137 | 0.984 | 0.046 |
| **Optimal dose of beta-blockers** | 1.433 | 0.584 | 3.516 | 0.032 |
| **Optimal-dose HF quadruple therapy** | 1.682 | 0.862 | 2.145 | 0.045 |

OR: odd ratio; CI: confidence interval

#### Correlation between the evolution of endothelial function and echography parameters

The improvement of endothelial function was associated with an increase in LVEF and cardiac output (R=0.339, p<0.001; R=-0.046, p=0.042, respectively). Additionally, the increase in the EQI was correlated with a decrease in the GLS (R=0.261, p=0.01).

#### Correlation between the evolution of endothelial function and hemodynamic parameters

The enhancement of endothelial function was associated with increased cardiac output (r=-0.226, p=0.007) and decreased systemic vascular resistance (r=0.269, p=0.03) (Supplementary Table 1).

**Supplementary Table 1:** Correlations between changes in hemodynamic parameters (Δ) and endothelial function.

| **Hemodynamic Parameter ($\Delta$)** | **Correlation Coefficient (R)** | **p-value** |
| --- | --- | --- |
| **Stroke Volume** | -0.065 | 0.61 |
| **Cardiac Output** | -0.226 | 0.007 |
| **Cardiac Index** | -0.188 | 0.134 |
| **Systemic Vascular Resistance** | 0.269 | 0.030 |
| **Systolic time ratio** | -0.155 | 0.217 |
| **Left Cardiac Work Index** | -0.186 | 0.139 |
| **Thoracic Fluid Content** | 0.057 | 0.65 |

$\Delta$ = change from baseline to 3 months: value at baseline – value at 3<sup>rd</sup> month

### One-year follow-up

The mean follow-up duration was 12 months. During this period, 5 out of 120 patients with HFrEF died, resulting in a survival rate of 93.2%.

Among the study population, 44 hospitalizations due to acute heart failure occurred one year after treatment, with a survival rate without major adverse cardiac events of 63.3%.

#### Endothelial function and prognosis

After one year of follow-up, patients with endothelial dysfunction at baseline had higher rates of mortality and rehospitalization compared to those with normal endothelial function (p < 0.001) (Supplementary Table 2).

**Supplementary Table 2.** The 12-month mortality and rehospitalizations according to baseline endothelial function in the study population.

| **Endothelial Function at Baseline** | **N** | **Deaths (n)** | **1-Year Survival Rate** | **Rehospitalizations (n)** | **Event-Free Survival Rate (1-Year)** |
| --- | --- | --- | --- | --- | --- |
| **Normal (EQI $\geq 2$)** | 26 | 0 | 100% | 5 | 80.8% |
| **Moderately Impaired ($1 \leq \text{EQI} < 2$)** | 43 | 1 | 97.7% | 12 | 72.1% |
| **Impaired (EQI $< 1$)** | 51 | 4 | 92.1% | 27 | 47.1% |

EQI: Endothelium Quality Index.

EQI: Endothelium Quality Index.

Improvement in endothelial function under optimized medical therapy was significantly associated with reduced mortality in our cohort. ROC curve analysis demonstrated an area under the curve of 0.82 (95% CI: 0.71–0.94, p = 0.01) (Supplementary Figure 1). A change in the EQI of less than 0.2 was identified as a predictive threshold for one-year mortality, with a sensitivity of 100% and a specificity of 47%.

Similarly, improvement in endothelial function was significantly associated with a reduction in rehospitalization rate. ROC curve analysis revealed an area under the curve of 0.837 (95% CI: 0.755– 0.919, p < 0.001). Our study found that a Δ EQI value below 0.2 predicted one-year rehospitalization for acute heart failure in patients with HFrEF, with a sensitivity of 88.6% and a specificity of 58% (Supplementary Figure 2).

## Discussion

The present study aimed to explore endothelial function in patients with chronic HFrEF and its association with echography and hemodynamic parameters following an optimized treatment. Furthermore, this study investigated the predictive role of endothelial function in the prognosis of chronic HFrEF.

The primary finding of the present study revealed that patients with chronic HFrEF commonly exhibit endothelial dysfunction. Moreover, improvements in endothelial function following optimized medical treatment were associated with favorable changes in echocardiographic and hemodynamic parameters. Interestingly, endothelial dysfunction emerged as a potential long-term predictor of major adverse cardiac events.

The findings of the present study demonstrated that 78.3% of patients with chronic HFrEF experienced endothelial dysfunction with a mean EQI of 1.27±0.6. In the literature, the assessment of endothelial function in chronic HFrEF was not well documented. Notably, a most recent meta-analysis demonstrated that patients with CHF with preserved ejection fraction exhibited endothelial dysfunction (12). Our results confirmed that endothelial dysfunction plays a crucial role in the pathogenesis and development of heart failure (13).

Endothelial dysfunction is characterized by a decrease in the bioavailability of NO, which becomes particularly relevant in coronary vessels, can lead to a reduction in myocardial perfusion, coronary flow, and impaired ventricular function (14). Therefore, endothelial dysfunction is associated with increased vascular stiffness, contributing to myocardial damage and a decline in cardiac performance (15).

The impairment in endothelial function can occur not only in the presence of coronary artery disease but also in other conditions such as diabetes, hypertension, and chronic kidney disease, underscoring the critical role of impaired endothelial vasodilation in the pathogenesis of heart failure (16). Endothelial dysfunction is determined by numerous alterations, such as the activation of the sympathetic nervous system and the increase in a systemic pro-inflammatory state associated with the release of circulating cytokines that downregulate eNOS expression and aggravate skeletal muscle atrophy, leading to exercise capacity in heart failure patients (17). On the other hand, the reduced expression of eNOS associated with a low production of NO may be elicited by the exacerbation of oxidative stress (18). The alteration of the redox state in CHF is a mechanism responsible for the decline of cardiac performance, initiated through direct damage caused by myocardial ischemia (19). In CHF, excessive oxidative stress is attributed to impaired endogenous antioxidant defense mechanisms, resulting from eNOS uncoupling and disrupted NO production (20). This redox imbalance contributes to dysfunctional calcium handling during systole and compromises myocardial relaxation (21). Furthermore, oxidative modifications increase endothelial permeability and compromise the structural integrity of the endothelium, exacerbating vascular dysfunction (21). In CHF patients, endothelin-1 (ET) is excessively upregulated, but NO is upregulated at a slower rate than ET, which often shows that the NO/ET balance shifts to ET. Elevated levels of ET contribute to increased vascular resistance, promote vascular smooth muscle cell proliferation, and stimulate extracellular matrix production (22). These alterations lead to excessive synthesis and accumulation of structural matrix proteins, resulting in myocardial fibrosis, increased myocardial stiffness, and diastolic dysfunction (23).

Consequently, endothelial dysfunction was associated with impaired cardiac function, as evidenced by a reduced LVEF with a mean of 29.92 ± 7.06%, a diminished GLS (10.8% ± 2.7), and alterations in key hemodynamic parameters.

The findings of this study revealed that optimized treatment of heart failure that gathers renin-angiotensin-aldosterone system inhibitors (i.e., Sacubitril and Valsartan), beta-blockers, aldosterone antagonists, and SGLT2 was effective in the enhancement of endothelial function from 1.27 ± 0.62 to 1.44 ± 0.67 (p<0.001).

The present study revealed that treatment with Sacubitril and Valsartan was a predictive factor for improving endothelial function. Our results were comparable with those of a study among patients with HFrEF and dilated cardiomyopathy, demonstrating that Sacubitril and Valsartan were effective in enhancing endothelial function, diastolic function, and mitral regurgitation (24). Interestingly, a 6-month administration of Sacubitril and Valsartan is safe and exerts beneficial effects on arterial stiffness, oxidative stress, platelet aggregation, and inflammatory biomarkers, which are critically involved in the pathophysiology of endothelial dysfunction and heart failure (25).

Beyond Sacubitril and Valsartan, our findings also highlighted the role of beta-blockers as independent predictors of improved endothelial function. Our results were in line with a meta-analysis showing that the flow-mediated dilation was enhanced with beta-blocker administration (26). Furthermore, beta-blockers were effective in improving the expression of pro-inflammatory biomarkers and heart failure symptoms (27). The beneficial effects of beta-blockers may be attributed not only to their vasodilatory properties but also to their antioxidant, antiproliferative, anti-hypertrophic, angiogenic, and anti-apoptotic effects (28). Consequently, beta-blockers that exert vasodilatory effects through NO pathways could represent a promising therapeutic strategy for the restoration of endothelial function (26).

Angiotensin-converting enzyme inhibitors have also demonstrated significant benefits in patients with HFrEF. These agents are well established for their antihypertensive properties and their ability to delay the progression of left ventricular remodeling (29). In our study, treatment with Angiotensin-converting enzyme inhibitors was associated with improved endothelial function, supporting findings from previous literature. Notably, a meta-analysis showed that Angiotensin-converting enzyme inhibitors significantly enhance flow-mediated dilation in patients with endothelial dysfunction. Moreover, their efficacy was superior to calcium channel blockers and beta-blockers (30). Our findings are further supported by a recent study in which 12-month therapy with perindopril in patients with HFrEF led to notable improvements in both macrovascular and microvascular endothelial function, as measured by photoplethysmography (31). By contrast, although angiotensin II receptor blockers are also used in the treatment of heart failure and have shown cardioprotective effects, their direct impact on endothelial function is less well-established (32).

The fourth class of optimized treatment was SGLT2, which emerged in our study as a significant predictive factor for improved endothelial function in patients with chronic heart failure. Our finding was in line with a 2023 study that highlighted the ability of SGLT to enhance flow-mediated dilation in individuals with elevated cardiovascular risk (33). SGLT2 primarily act on the proximal renal tubule to reduce glucose and sodium reabsorption, thereby inducing glycosuria, natriuresis, and diuresis (34). This reduction in intracellular sodium has been proposed to improve calcium handling in cardiomyocytes, contributing to enhanced cardiac contractility. Moreover, these agents exhibit additional pleiotropic effects, including the reduction of oxidative stress, myocardial fibrosis, inflammation, and the promotion of autophagy (34). Specifically, empagliflozin has been shown to modulate key molecular pathways by enhancing eNOS-dependent protein kinase G Iα oxidation and downregulating inflammatory markers.

Our study demonstrated that the improvement in endothelial function under medical treatment was significantly correlated with echocardiographic parameters, including LVEF, GLS, and subaortic velocity time integral, as well as hemodynamic parameters assessed by IC, such as cardiac output and vascular resistance. These findings suggest that medical therapy may have a measurable impact on circulatory dynamics, indicating that treatment can not only enhance endothelial function but also positively influence hemodynamic measurements.

In the present study, we recorded a total of 49 major adverse events, including 5 deaths and 44 rehospitalizations due to acute decompensated heart failure, with a cumulative survival rate of 93.2% over a one-year follow-up. Particularly, a higher mortality rate was recorded in the group of patients with impaired endothelial function at baseline (4 deaths), compared to those with moderately impaired function (1 death), while no deaths were recorded in the group with normal endothelial function.

Regarding hospital readmissions, our study showed that patients with severely impaired endothelial function at baseline had a significantly higher rate of readmissions (27 readmissions) compared to those with normal or moderately impaired endothelial function (p = 0.008). These findings are consistent with a study among CHF, revealing that patients with low flow-mediation dilation experienced adverse cardiovascular events, compared to patients in the high flow-mediation dilation group (p < 0.01) (35). Similarly, a study among 242 patients with chronic heart failure demonstrated that post-hyperemic forearm blood flow was identified as an independent predictor of major events (i.e., death, myocardial infarction, angina, stroke, or hospitalization due to heart failure) in CHF patients during a 5-year of follow-up (36).

Interestingly, our findings revealed that the improvement in endothelial function with an increase of EQI of 0.2 was associated with a decrease in hospitalization and mortality rate, with an area under curve of 0.82 and 0.837, respectively.

Our findings reinforce the hypothesis that endothelial dysfunction is not only a key pathophysiological feature of heart failure but also a significant prognostic marker. Numerous studies have shown that it is independently associated with increased risks of all-cause mortality, non-fatal myocardial infarction, and hospitalizations due to heart failure decompensation (37).

This adverse prognostic impact appears to be especially prominent in ischemic heart failure, where impaired endothelium-dependent vasodilation plays a central pathophysiological role (35).

Our findings underscore the role of endothelial function as an excellent surrogate marker in clinical settings, capable of reflecting both disease severity and therapeutic response. Incorporating routine endothelial function assessment into heart failure management may thus offer clinicians a practical and non-invasive tool for risk stratification and treatment monitoring.

## Study limitations

The present study has several limitations that should be acknowledged. First, the sample size remains relatively small for generalizing the findings to the broader population of patients with HFrEF. A larger cohort would strengthen the statistical power and enhance the external validity of the results. Second, the monocentric nature of the study may limit the applicability of the findings to other clinical settings, given potential regional differences in patient management, treatment protocols, and demographic characteristics. Additionally, the follow-up period of one year, although sufficient to capture short-term effects of optimized medical therapy, may not fully reflect the long-term trajectory of endothelial function, hemodynamic changes, rehospitalization rates, and mortality.

## Conclusion

Patients with HFrEF commonly tend to exhibit endothelial dysfunction. Interestingly, optimized guideline-directed medical therapy significantly improved endothelial function over three months. The vascular improvement was closely linked to favorable changes in hemodynamic parameters and echocardiographic measures, including left ventricular function and strain. Importantly, we identified endothelial dysfunction not only as a frequent finding in HFrEF but also as a powerful predictive marker of one-year outcomes. Patients with persistent endothelial dysfunction had higher rates of mortality and rehospitalization, whereas those showing significant improvement in endothelial function experienced better clinical trajectories. This association underscores the potential of endothelial function as a dynamic, non-invasive marker to monitor treatment response and stratify risk.

## Data availability

The data that support the findings of this study are available from the corresponding author upon reasonable request.

## Acknowledgments

All authors would like to thank all the volunteers who participated in the present study.

## Authors contribution

**SC** Conceptualization, methodology, investigation, formal analysis, resources, writing-original draft; **MAH** conceptualization, formal analysis, writing-original draft, resources; **NL** data curation, investigation, methodology, writing-original draft; **MJ** investigation, resources; **AB** investigation, resources; **MJ** investigation, data curation, resources; **HG** investigation, resources; **AG** investigation, data curation; **ZT** investigation, data curation; **TE** investigation, resources; **FT** investigation, data curation, validation; **RG** investigation, data curation, validation; **LA** investigation, methodology, data curation, validation, supervision

All authors have read and approved the final version of the manuscript and agree with the order of presentation of the authors.

## Abbreviations

CHF: chronic heart failure
ET: endothelin-1
EQI: Endothelium Quality Index
GLS: Global longitudinal strain
HFrEF: heart failure with reduced ejection fraction
IC: impedance cardiography
LVEF: left ventricular ejection fraction
NO: nitric oxide
ROC: Receiver Operating Characteristic
SGLT2: sodium-glucose co-transporter 2 inhibitors

## Figure legends

**Supplementary Figure 1.**
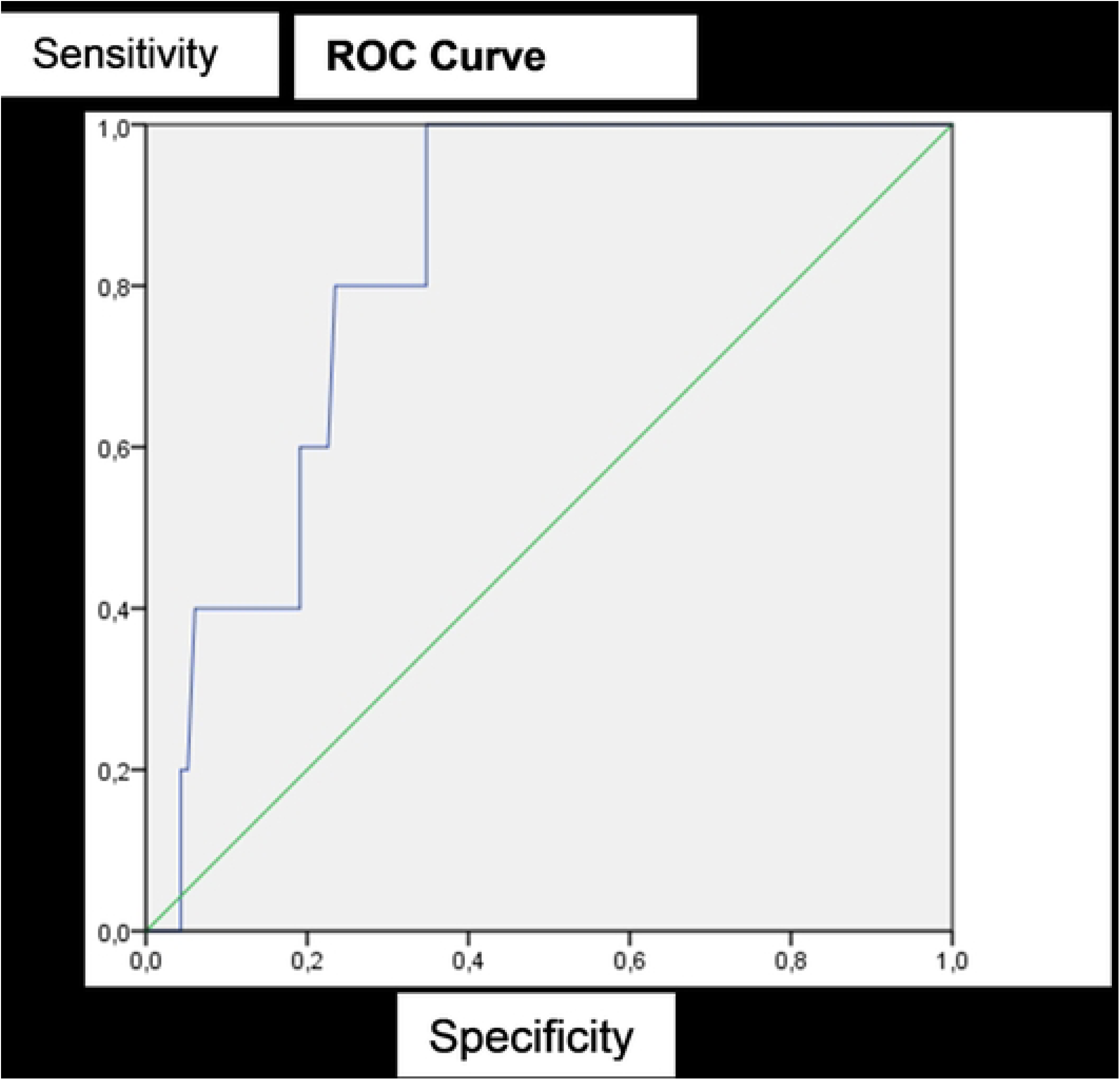
Receiver operating characteristic curve highlighting the association between one-year mortality and changes in endothelial function under optimized therapy

**Supplementary Figure 2.**
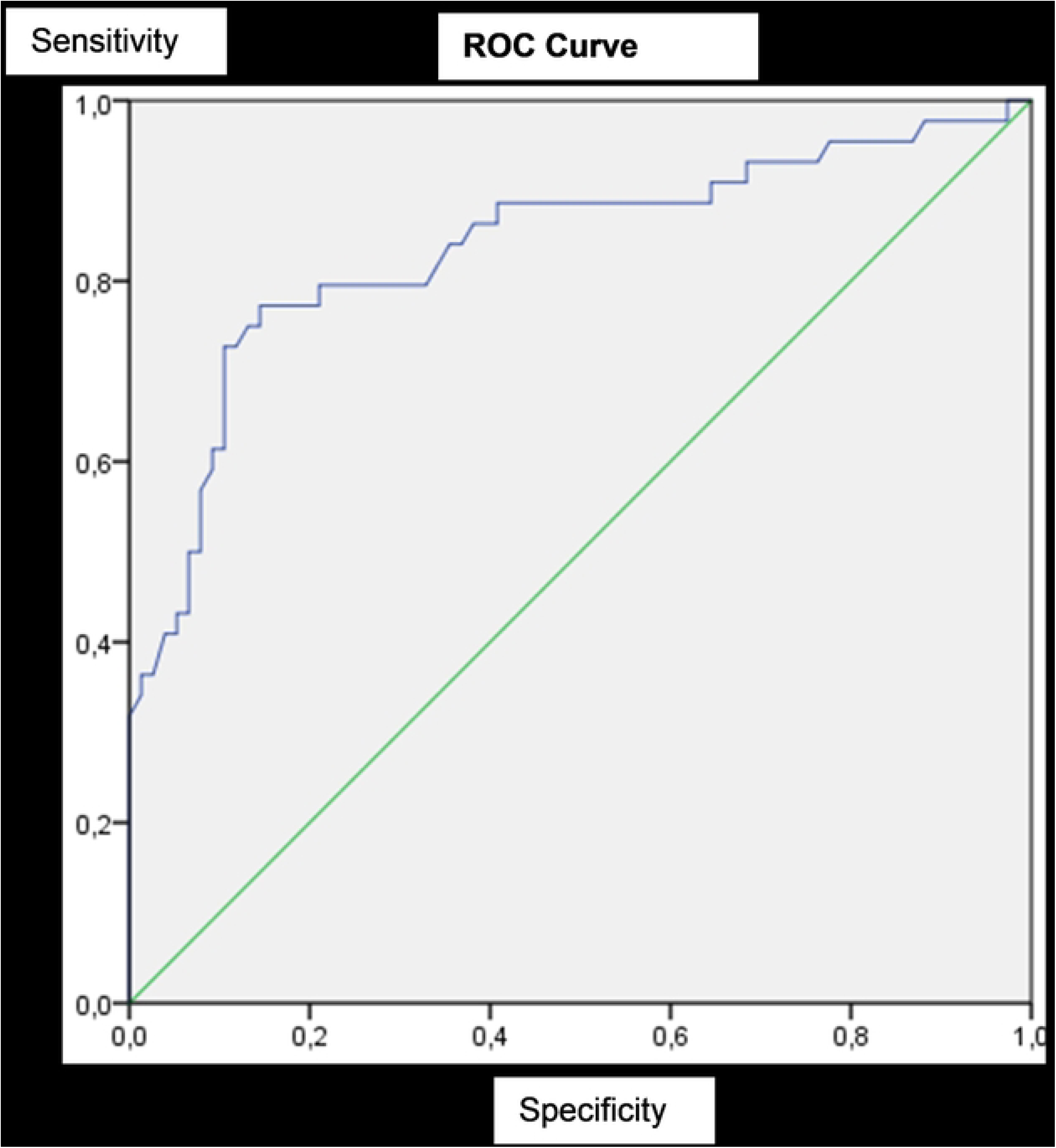
The association between 1-year rehospitalizations due to acute heart failure and changes in endothelial function under optimized therapy.

